# Flexible top-down modulation in human ventral temporal cortex

**DOI:** 10.1101/279935

**Authors:** Ruyuan Zhang, Kendrick Kay

## Abstract

Visual neuroscientists have long characterized attention as inducing a scaling or additive effect on fixed parametric functions describing neural responses (e.g., contrast response functions). Here, we instead propose that top-down effects are more complex and manifest in ways that depend not only on attention but also other cognitive processes involved in executing a task. To substantiate this theory, we analyze fMRI responses in human ventral temporal cortex (VTC) in a study where stimulus eccentricity and cognitive task are varied. We find that as stimuli are presented farther into the periphery, bottom-up stimulus-driven responses decline but top-down attentional enhancement increases substantially. This disproportionate enhancement of weak responses cannot be easily explained by conventional models of attention. Furthermore, we find that attentional effects depend on the specific cognitive task performed by the subject, indicating the influence of additional cognitive processes other than attention (e.g., decision-making). The effects we observe replicate in an independent experiment from the same study, and also generalize to a separate study involving different stimulus manipulations (contrast and phase coherence). Our results suggest that a quantitative understanding of top-down modulation requires more nuanced and more precise characterization of multiple cognitive factors involved in completing a perceptual task.

## INTRODUCTION

To tackle the immense size and complexity of visual inputs, the brain concentrates limited attentional resources on the most informative aspects of visual inputs. The mechanisms of attentional allocation have been an active research area in past years, because of the pivotal role that attention plays in different sensory processes, such as feature binding (Treisman AM and G Gelade 1980), object recognition (Walther D et al. 2002), and scene understanding (Itti L et al. 1998). Neuroscientists are particularly interested in the neural substrates of attention. Converging evidence from primate electrophysiology and human neuroimaging suggests that attention induces enhancement in microscopic neuronal activity (Reynolds JH et al. 2000) as well as macroscopic cortical responses (Gandhi SP et al. 1999; Murray SO and E Wojciulik 2004). Such attention-induced response enhancement is thought to produce more robust sensory representations (Kastner S and LG Ungerleider 2000; Reynolds JH and L Chelazzi 2004).

Despite the well-established finding of attentional enhancement of neural responses, the precise quantitative nature of attentional enhancement remains unclear. One conventional approach to tackling this issue is to characterize the impact of attention on the shape of contrast response functions (CRFs) (Reynolds JH *et al.* 2000; Buracas GT and GM Boynton 2007; Boynton GM 2009), that is, functions describing the relationship between input stimulus contrast and output neural response. Under the assumption that neural responses follow a fixed parametric form (such as the commonly used Naka-Rushton function (Albrecht DG and DB Hamilton 1982)), attention is characterized as imposing a scaling or additive effect on either input contrast or output response. As illustrated in Figure 1, attention could have the effect of amplifying the overall CRF (Figure 1A), enhancing the input contrast (Figure 1B), or inducing a baseline shift (Figure 1C). Though mathematically elegant, this approach cannot fully explain some experimental measurements found in the attention literature (Luck SJ et al. 1997; Reynolds JH *et al.* 2000; Li X et al. 2008; Murray SO 2008), and moreover, it is not clear whether this *fixed-parameter approach* generalizes to stimulus dimensions other than contrast. Thus, it remains an open question whether the approach provides a satisfactory account of attentional effects.

**Figure 1.**
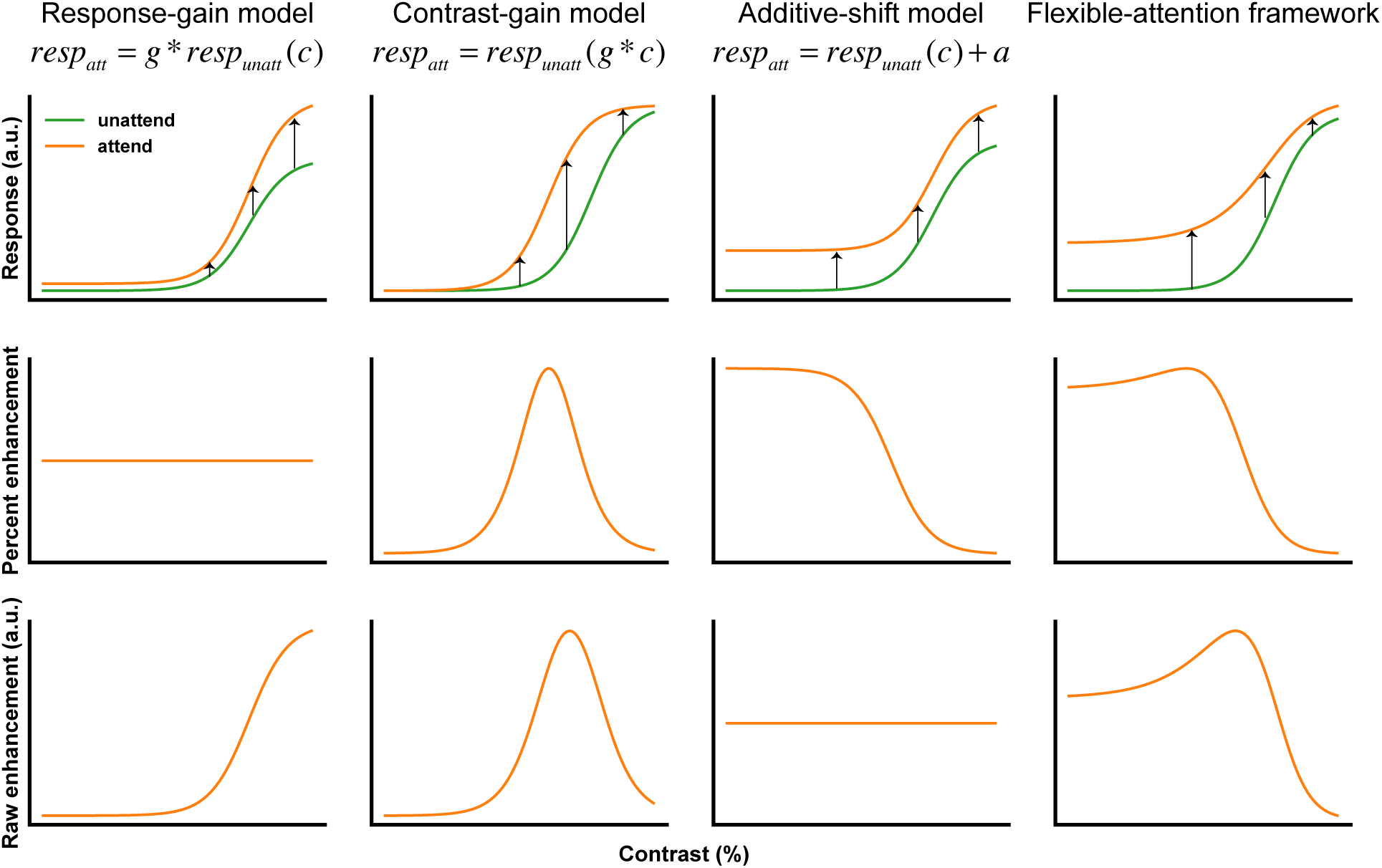
Schematics of conventional models of attention and the flexible-attention framework. The first row depicts contrast response functions under unattended (*resp*_*unatt*_) and attended (*resp*_*att*_) conditions. Arrows indicate attentional enhancement. The second and third rows depict the amount of attentional enhancement under two different metrics: percent enhancement (Equation 1) and the raw enhancement (Equation 2), respectively. The *response-gain model* posits that attention imposes a scaling effect (*g*) on the output, and therefore predicts that percent enhancement is a flat function of contrast. The *contrast-gain model* posits that attention imposes a scaling effect (*g*) on the input contrast, and predicts that both percent enhancement and raw enhancement are inverted U-shaped functions. The *additive-shift model* posits that attention imposes an additive effect (*a*) on the output, and predicts that raw enhancement is a flat function of contrast. In contrast to these fixed-parameter approaches, the *flexible-attention framework* allows for the possibility that attentional effects are neither constant in percent enhancement nor constant in raw enhancement. Here we depict one possibility where attention disproportionately enhances low-contrast responses.

In this paper, we advocate moving beyond the fixed-parameter approach and argue that it is more appropriate to consider attention as a flexible process that depends on the specific stimuli and task demands faced by the observer. In this *flexible-attention framework*, attention is not a simple binary variable (i.e., ‘present’, ‘absent’), but rather, attentional effects depend on specific properties of the cognitive processes involved in a task (e.g., whether a detection or a discrimination task is being performed). Since tasks are remarkably diverse, the effects of attention on neural responses may manifest in different ways, and a fixed parametric function might not accurately capture attentional effects observed in an arbitrary experiment. Empirical evidence inspiring the flexible-attention framework comes from a recent study (Kay KN and JD Yeatman 2017) in which we measured cortical responses to different stimulus categories while subjects performed different tasks (henceforth referred to as the *category study*).

Here, we strengthen support for the flexible-attention framework through a re-examination of experimental measurements from an independent study (Kay KN et al. 2015). In this study, cortical responses were measured for different stimulus positions while subjects performed different tasks (henceforth referred to as the *position study*). We quantify attentional effects in human ventral temporal cortex (VTC) as a function of stimulus eccentricity, and apply the same type of analysis to the category study, thereby allowing direct comparison of results. Across studies, we show that weak stimulus-driven responses receive disproportionately large attentional enhancements and attentional enhancements are more pronounced for certain tasks compared to others. Such effects are not well explained by conventional models of attention, and therefore suggest the need to develop a more flexible framework for attention. In the Discussion, we propose specific ways in which the concept of “flexible attention” might be formalized into a quantitative model.

## MATERIALS AND METHODS

### Experiment and MRI data acquisition

Three adults participated in the position study (Kay KN *et al.* (2015)). In the *task experiment* (Figure 2), face stimuli (3.2° diameter) appeared at different positions of a 5 × 5 spatial grid (1.5° spacing). This grid sampled six distinct eccentricities (0°, 1.5°, 2.1°, 3°, 3.4° and 4.2°). Each trial consisted of 7 sequentially presented faces (500ms/face) at a single position but with various identities and viewpoints. Some trials involved two consecutive faces sharing the same identity but different viewpoints, and some trials involved a red dot appearing at the center of the faces (coincident with one of the 7 faces). A stream of digits (0.3° x 0.3°) was placed at the center-of-gaze. In a given run, participants were instructed to perform either (1) a digit task, during which participants pressed a button whenever the same digit repeated; (2) a dot task, during which participants pressed a button whenever a red dot appeared; or (3) a face task, during which participants pressed a button whenever the same face identity repeated within a trial. Participants fixated the central stream of digits during all three tasks (verified using an eyetracker). There were 75 experimental conditions (25 locations x 3 tasks) and 8 trials for each condition over the course of the experiment. All experimental details are described in Kay KN *et al.* (2015).

**Figure 2.**
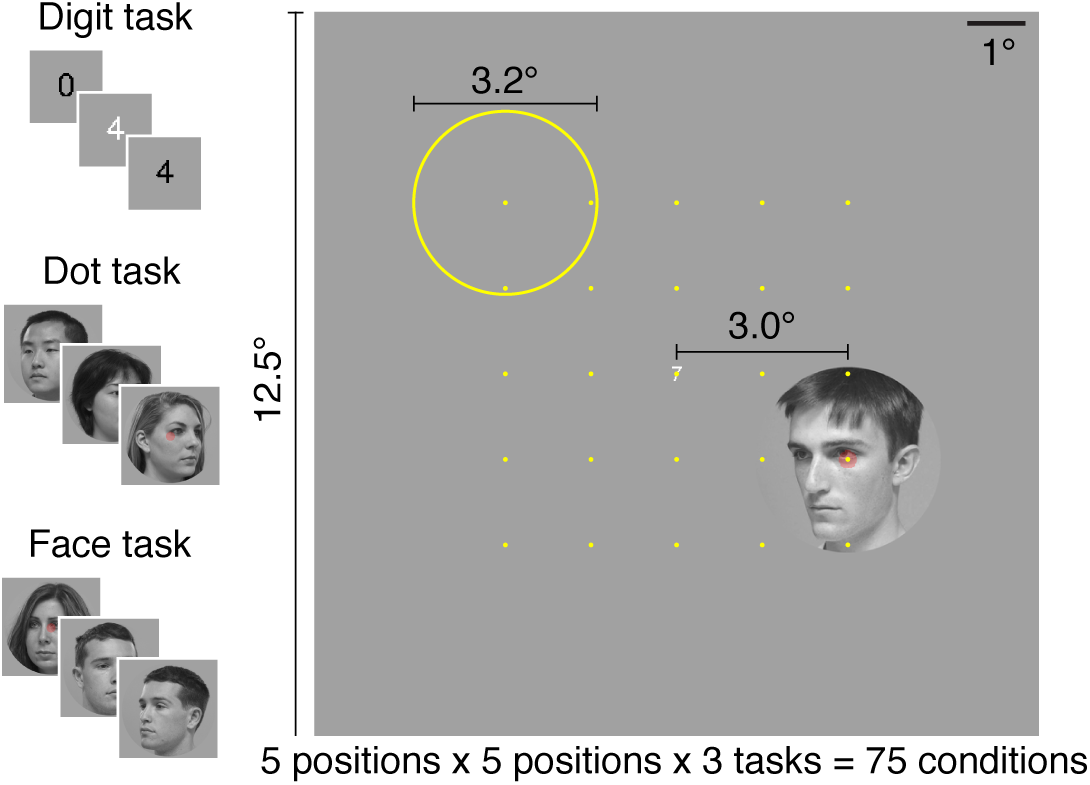
Stimuli and tasks from the position study (Kay KN *et al.* (2015)). In a given trial, a sequence of face stimuli (7 face images) appears in one of twenty-five positions. The *digit task* is a one-back task on the stream of digits at the center-of-gaze. The *dot task* is to detect the occurrence of a red dot on the faces. The *face task* is a one-back task on the identity of the faces. Subjects maintained central fixation, and stimuli were identical across the three tasks.

The position study included another experiment, called the *interleaved-task experiment*. This experiment was the same as the task experiment (Figure 2) except that the three tasks were randomly intermixed in a trial-by-trial fashion within each run. A central red letter (0.3° x 0.3°) presented at the beginning of each trial served as a cue for which task to perform. This experiment provides an additional, independent set of data.

Functional MRI data were collected at the Stanford Center for Cognitive and Neurobiological Imaging using a 3T GE Signa MR750 scanner, a Nova 16-channel visual RF coil, and a gradient-echo EPI pulse sequence (TR 2 s, 2-mm voxels). The fMRI data were pre-processed by performing slice time correction, spatial distortion correction and motion correction. The fMRI data were further analyzed using GLMdenoise (Kay KN et al. 2013) to estimate the percent BOLD signal change (beta weight) evoked by each stimulus location under each task. This analysis also generated 100 bootstrap samples of beta weights via resampling of scanning runs.

Visual field maps (V1, V2, V3, and hV4) were defined using standard retinotopic mapping scans. Three face-selective regions (inferior occipital gyrus, IOG-faces/OFA (abbreviated IOG); posterior fusiform gyrus, pFus-faces/FFA-1 (abbreviated pFus); and middle fusiform gyrus, mFus-faces/FFA-2 (abbreviated mFus)) were defined using independent functional localizer scans. We also defined IPS as an additional ROI (beyond that described the original paper). Specifically, we used the IPS-0 region from an atlas of visual topographic organization (Wang L et al. 2015); this choice is reasonable given the limited coverage of parietal cortex available in the position study and the localization of top-down modulation to IPS-0/1 as shown in (Kay KN and JD Yeatman 2017).

### Region-level analysis

After the GLM analysis, we pooled voxels within corresponding regions of interests (ROIs) across subjects and hemispheres. The same voxel selection criterion (goodness-of-fit of the population receptive field model) used in our previous paper was applied to exclude non-spatially selective voxels (Kay KN *et al.* 2015). To calculate region-level responses, we first computed the median across bootstrap samples to obtain the response of each voxel to the 75 experimental conditions. The responses of individual voxels were then positively rectified to remove negative responses. Finally, we calculated the region-level response by computing the mean across voxels.

Two metrics were used to quantify the magnitude of attentional effects: *percent enhancement* and *raw enhancement*, which are defined as follows:

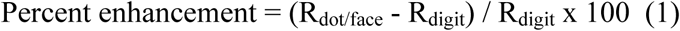

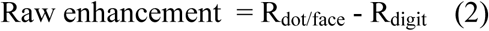

where R_dot/face_ indicates an ROI’s response for a stimulus location in the dot or the face task and R_digit_ indicates the ROI’s response for the same location in the digit task. This calculation provides 50 values (25 for the dot task and 25 for the face task) for each metric.

### Analysis of data from the category study

We reanalyzed data from the category study (Kay KN and JD Yeatman 2017) using the same methods described above for the position study. In brief, the category study involved presentation of words, faces, and other stimulus categories varying in contrast and phase coherence. Subjects performed one of three tasks: (1) a fixation task, during which participants pressed a button whenever the fixation dot turned red; (2) a categorization task, during which participants reported whether the stimulus was a word, face, or neither; and (3) a one-back task, during which participants pressed a button whenever an image was repeated twice in a row.

In Figures 6–8, we directly compare results across the position and category studies. To facilitate comparison, we pooled voxels from pFus and mFus in the position study to match the definition of fusiform face area (FFA) in the category study. Also, since overall response amplitudes might vary for incidental reasons across subjects, we normalized bottom-up responses (responses during the digit task of the position study and responses during the fixation task of the category study) by dividing by the maximal response amplitude observed in each study and ROI. For example, the full set of responses measured from FFA in the category study (including both contrast and phase coherence conditions) was divided by the maximum response. Note that this normalization affects raw enhancement values but not percent enhancement values.

### Error bars

Unless otherwise indicated, error bars indicate 68% confidence intervals, obtained by bootstrapping across locations that share the same eccentricity (position study) or bootstrapping across subjects (category study).

## RESULTS

### Cortical responses as a function of stimulus eccentricity and behavioral task

We refer to the main experiment in the position study as the *task experiment* (see Methods for details). In the task experiment, participants performed three different cognitive tasks on face stimuli that appeared at six different eccentricities while blood oxygenation level dependent (BOLD) signals in human ventral temporal cortex (VTC) were measured. Using face stimuli rather than artificial visual stimuli (e.g., checkerboards) produces strong responses not only in early visual areas but also in high-level category-selective regions. This allows us to assess attentional effects throughout the visual cortical hierarchy.

Participants performed three different tasks. The *digit task* is a one-back task on a stream of digits placed at the center-of-gaze. Face stimuli in this task are irrelevant to the participants, and the purpose of this task is to maintain participants’ attention at the central fixation point. Although participants may occasionally attend to the face stimuli, we interpret responses in the digit task as primarily reflecting bottom-up visual processing with minimal top-down influences. The *dot task* requires participants to detect the occasional appearance of a red dot superimposed on the face stimuli. In this task, face features (e.g., identity, viewpoint) are irrelevant to the participants. The *face task* requires participants to perform a one-back task on face identity; thus, face features in this task are highly relevant to the participants.

We summarized the responses of each region-of-interest (ROI) as a function of stimulus eccentricity, producing eccentricity-response functions (ERFs). This is analogous to conventional contrast-response functions where responses are plotted as a function of stimulus contrast. Examining the ERFs allows us to inspect whether attentional effects observed for contrast response functions generalize to other feature dimensions. We discovered several prominent effects. First, the evoked responses in high-level face-selective areas generally decrease as stimulus eccentricity increases (Figure 3), indicating that stimulus eccentricity, like contrast, has a strong influence on cortical responses. Second, the fact that responses increase from the dot task to the face task suggests that the brain enhances responses if the task requires detailed processing of the attended stimulus. Finally, the effect of task on cortical responses progressively developed along the visual cortical hierarchy, suggesting that attentional effects are more pronounced in brain regions whose representations are critical to successful execution of the task (i.e., face-selective regions for judging face identity).

**Figure 3.**
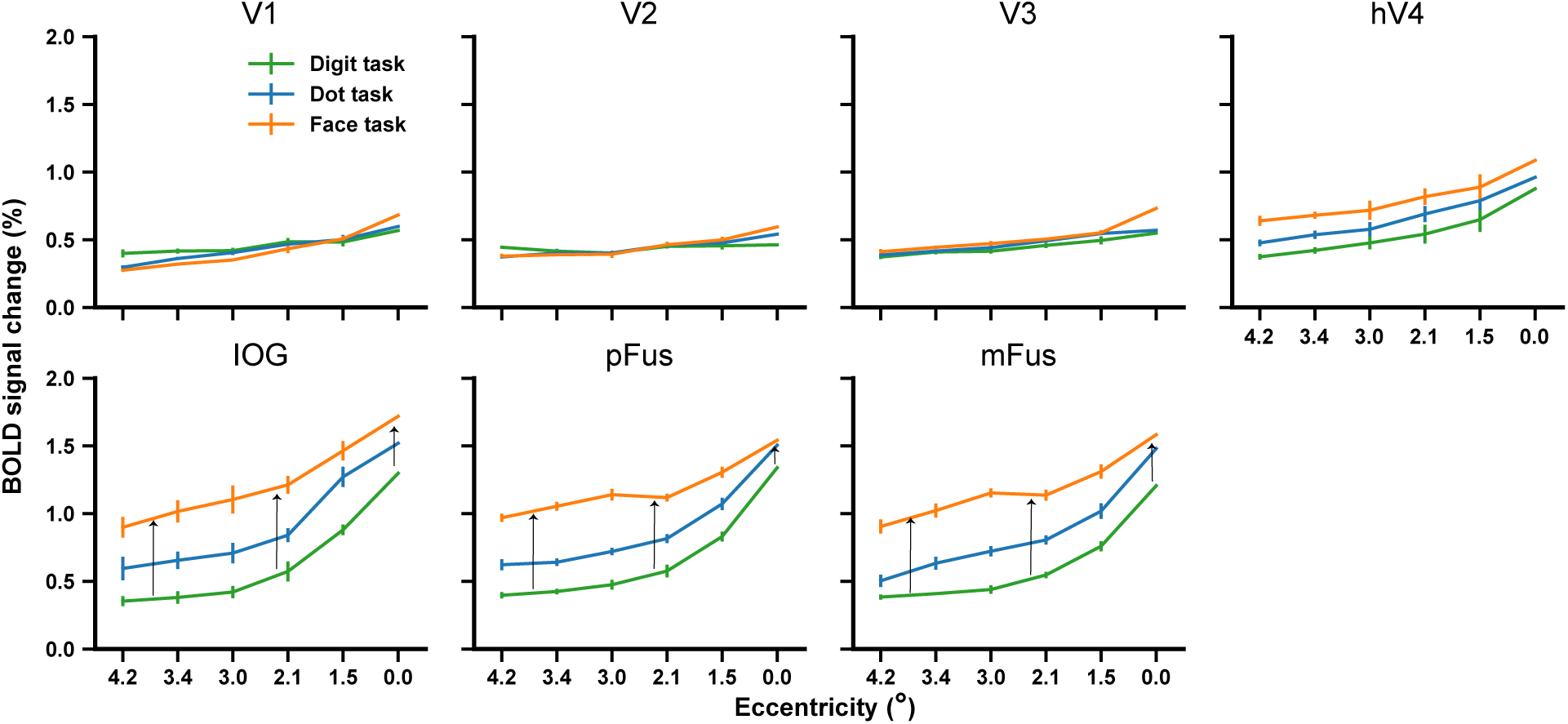
Percent BOLD signal change as a function of stimulus eccentricity and task. The order of stimulus eccentricity is reversed to make eccentricity-response functions visually comparable to contrast-response functions. BOLD responses are pooled across subjects and hemispheres (see Methods). Error bars indicate 68% confidence intervals on the bootstrapped mean of responses across locations at the same eccentricity (note that 0° corresponds to only one location and thus has no error estimate). Unless specifically mentioned, the same error-bar convention is used in subsequent figures. Responses in high-level visual areas exhibit substantial dependence on both eccentricity and task. Black arrows highlight the disproportionate attentional enhancement at high eccentricities, reminiscent of the schematic of the flexible-attention framework in Figure 1.

### Conventional models of attention cannot fully account for observed attentional effects

We next evaluate the accuracy of different attentional models. We quantified attentional effects as a function of stimulus eccentricity and task using two metrics: *percent enhancement* (Equation 1) and *raw enhancement* (Equation 2). These metrics were used because they allow direct assessment of the accuracy of the response-gain and the additive-shift models of attention (Figure 1). Results indicate that previously proposed models of attention do not fully account for the data (Figure 4). The reasons are as follows.

**Figure 4.**
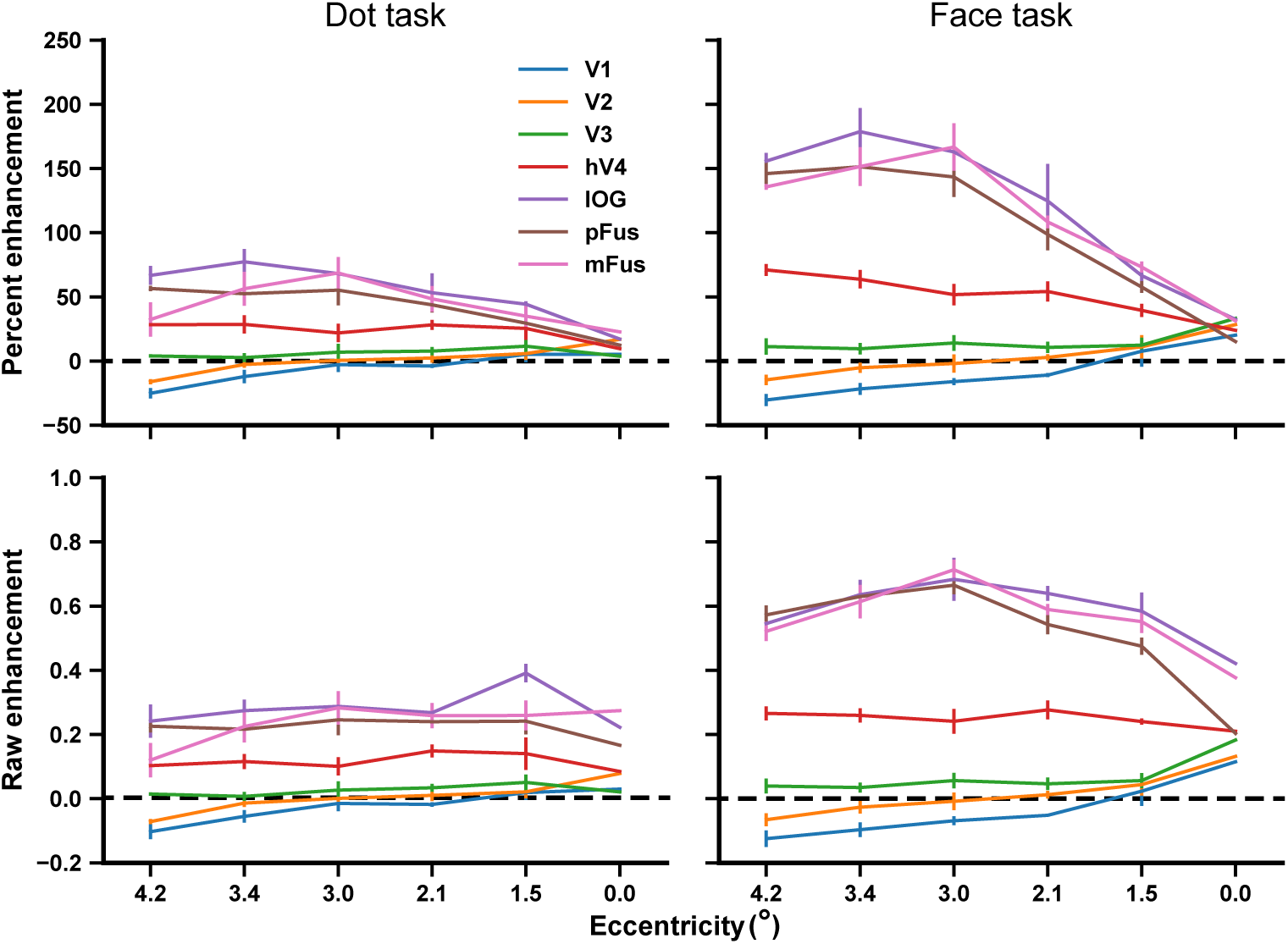
Attentional enhancement as a function of stimulus eccentricity and task. BOLD responses during the stimulus-relevant tasks (dot and face tasks) are expressed as percent enhancement (upper row) and raw enhancement (bottom row) relative to the responses during the digit task. The horizontal dashed line indicates no attentional enhancement. The magnitude of the attentional effect increases from fovea to periphery, from the dot task to the face task, and from low-level to high-level visual areas. This pattern is inconsistent with the three conventional models of attention (see explanations in main text). Note that the data point at 0° corresponds to only one location and thus has no error estimate.

First, the response-gain model posits that attention amplifies the overall magnitude of ERFs, leading to larger attentional effects when bottom-up stimulus-driven responses are larger, i.e., in the fovea. It also predicts percent enhancement will be a flat line as a function of stimulus eccentricity. These predictions are not consistent with Figure 4–in the face-selective regions, raw enhancement is not large in the fovea and there is a clear rising trend of percent enhancement from fovea to periphery.

Second, the additive-shift model posits that attention vertically shifts ERFs; thus, raw enhancement should be a flat function of stimulus eccentricity. This prediction seems only consistent with the data in the dot task. The dot task, however, involved no demands for processing face features and the ROIs exhibiting the largest attentional effects are face-selective regions (also see Discussion). In the face task, raw enhancement as a function of eccentricity is not a flat line and instead rises in face-selective regions as stimulus eccentricity increases.

Finally, the contrast-gain model predicts the largest percent enhancement and raw enhancement in middle levels of eccentricity, resulting in inverted U-shaped functions of percent enhancement and raw enhancement (Figure 1B). It is also unsuited here since the strongest attentional effects, under both metrics, appear in the far visual periphery (also see Discussion).

Since the results are inconsistent with attentional models proposed in previous literature, we propose the idea of flexible attention in which attentional effects do not necessarily conform to simple parametric changes. Before elaborating on this idea, we show first that the observed effects are not idiosyncratic features of this particular experiment but generalize across several stimulus and task manipulations.

### Reproducible effect of flexible attention on an independent dataset

All analyses thus far are based on the data from the task experiment where three different cognitive tasks were performed in different scanning runs. We also conducted an *interleaved-task* experiment in which tasks were interleaved in a trial-by-trial fashion within a run (see Methods for details). This experiment provides an independent dataset that can be used to confirm the observed effects. We applied the same analysis above on the data from the interleaved-task experiment. The two independent experiments yield highly consistent results (Figure 5), further supporting the presence of flexible attentional modulation.

**Figure 5.**
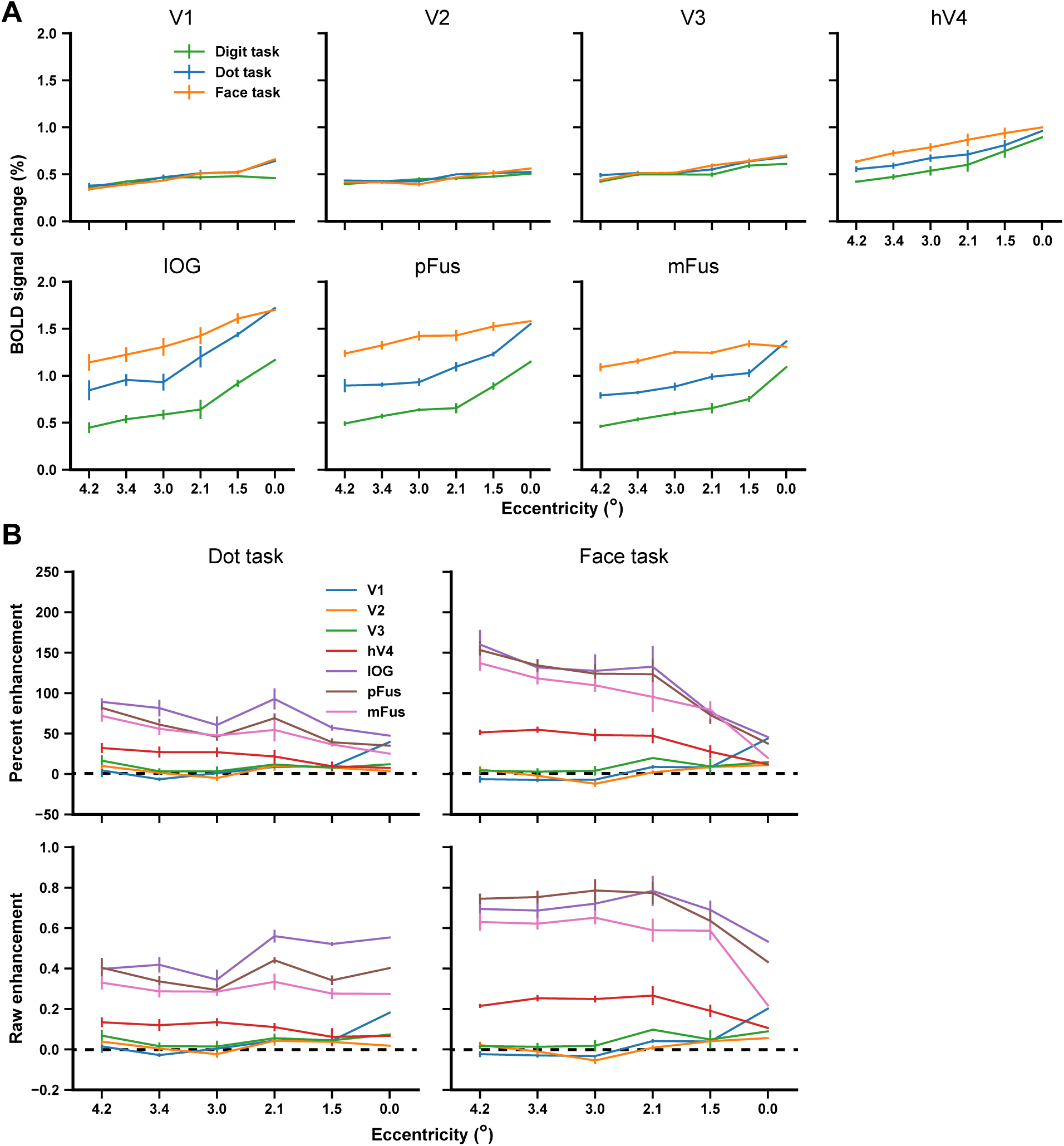
Disproportionate attentional enhancements at high eccentricities in the interleaved-task experiment. ***A-B***. Results are plotted in the same format as the results from the task experiment shown in Figures 3-4. Overall, the results from the two independent experiments are highly consistent.

### Evidence for flexible attention in other experimental manipulations

The form of attentional modulation discovered in this study, especially the dependency of attentional effects on the level of physical stimulus, has rarely been discussed in previous literature. However, we found similar effects in Kay KN and JD Yeatman (2017) (termed the *category study*) in which responses to different stimulus categories are investigated. In that study, we reported that attention selectively imposes larger scaling effects on weaker responses, a phenomenon termed “stimulus-specific scaling”. We thus consider applying the same analyses demonstrated above to the data from the category study. Exploiting the data from that study has two major attractions: (1) In the position study, only one stimulus feature–eccentricity–is manipulated. In the category study, stimuli are manipulated in both contrast and phase coherence, thus providing two extra feature dimensions that influence bottom-up visual processing. (2) The responses in another ROI–visual word form area (VWFA)–were also measured. This allows us to test whether our findings are specific to FFA or generalize to other high-level visual regions.

We extracted BOLD responses in FFA and VWFA toward their preferred stimulus categories–faces and words, respectively. To make data from the two studies more comparable, voxels from pFus and mFus in the position study were pooled, consistent with the definition of FFA in the category study. Furthermore, we highlight data from the stimulus-relevant tasks that yielded strongest attentional effects: the face task in the position study and the one-back task in the category study.

The two studies show a consistent pattern (Figure 6): attentional effects are larger for stimuli that evoke weak bottom-up responses (digit task in the position study and fixation task in the category study). As explained previously, neither the response-gain nor the additive-shift model of attention can account for the results. Instead, these results suggest the need for the flexible-attention framework (Figure 1D). One exception to the general pattern of large attentional enhancement at weak stimulus strength lies in phase coherence (Figure 6D, F). We speculate that this may be due to the fact that 0% phase coherence images contain pure noise on which it may be easier to perform a one-back decision (also see Discussion).

**Figure 6.**
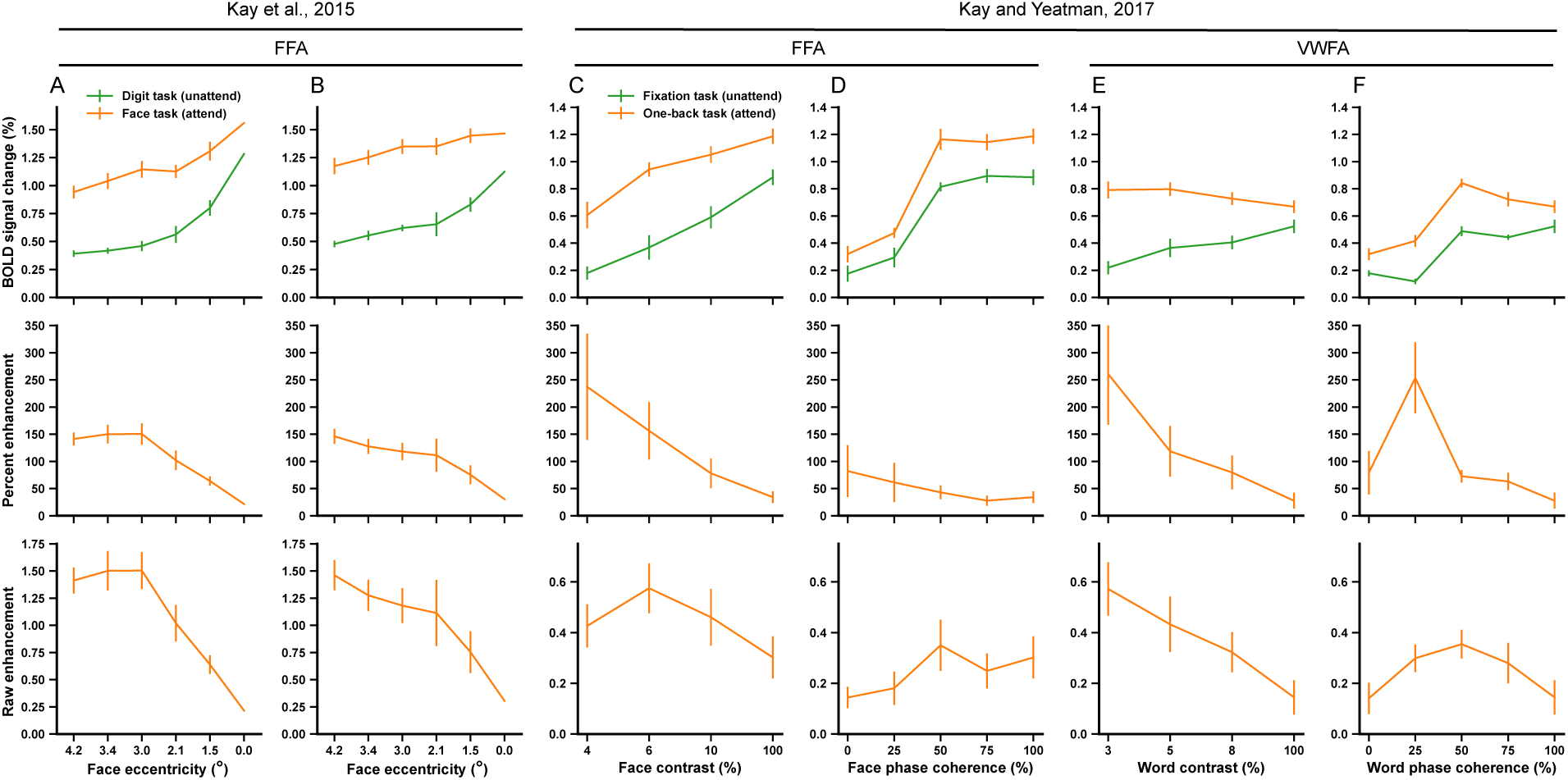
Disproportionate attentional enhancements generalize across experiments. Panels A–B show results from the position study for the task and the interleaved-task experiments, respectively. Panels C–F show results from the category study. Data from that study have been analyzed in the same way as Panels A–B, except that error bars reflect 68% confidence intervals on the mean across subjects (see Methods for details). Across metrics, the amount of attentional enhancement generally decreases as stimulus strength (eccentricity, contrast, phase coherence) increases. Enhancement tends to be greatest when stimulus strength is low and bottom-up responses (green curves in the first row) are weak.

To gain further insight into the relationship between bottom-up responses and the magnitude of attentional enhancement, we plot percent enhancement and raw enhancement values against the bottom-up responses across stimuli, tasks, studies, and ROIs (Figure 7). The clear inverse relationships between bottom-up responses and the amount of attentional effect indicate that attention disproportionately enhances weak neural responses.

**Figure 7.**
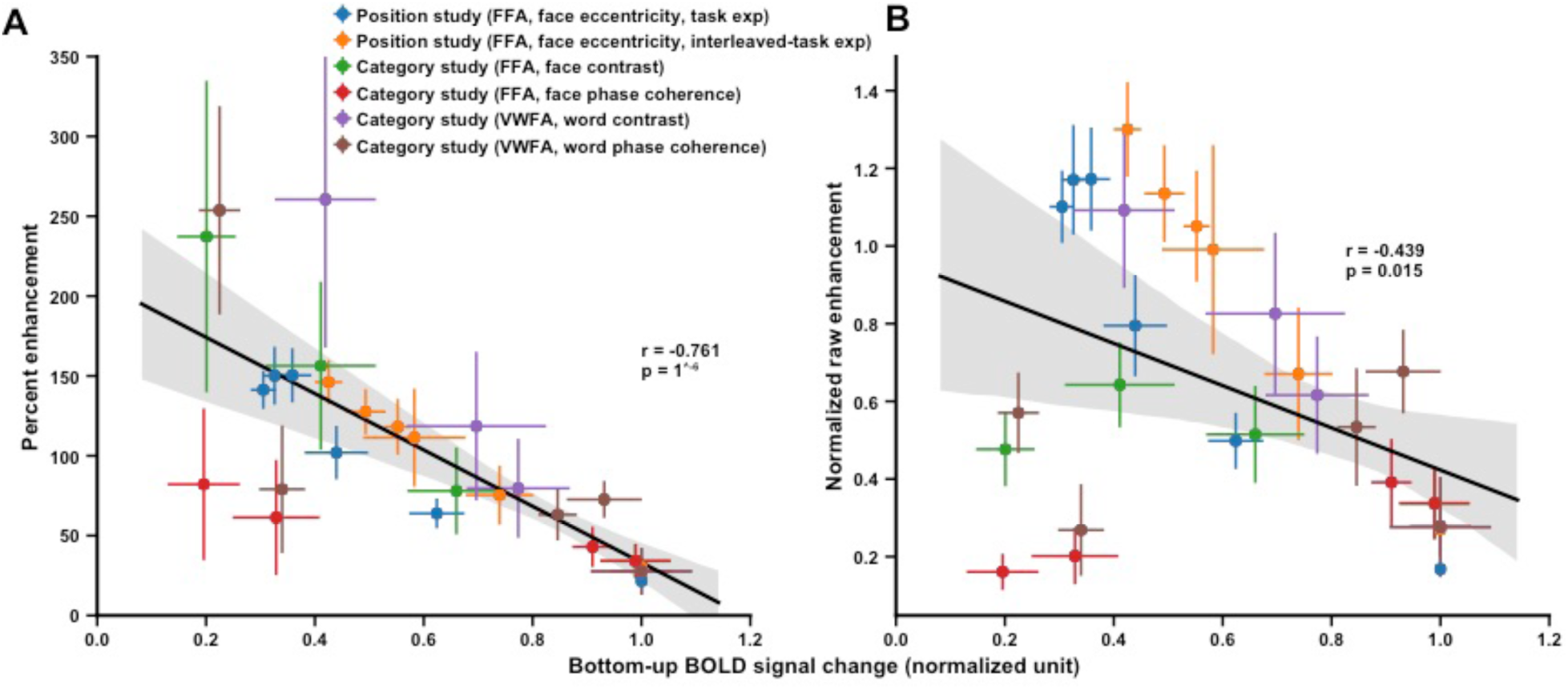
Inverse relationships between normalized bottom-up responses to percent enhancement (***A***), and normalized raw enhancement (***B***). All data points from the two studies depicted in Figure 6 are plotted. To ensure that BOLD responses from different ROIs and experiments are in comparable units, we normalize the full set of responses observed during the bottom-up tasks (the digit task in the position study and the fixation task in the category study) in each ROI to a maximum of 1 (see Methods). The shaded area indicates the 95% confidence interval of a bootstrapped best-fit line. The results demonstrate that attentional enhancement tends to be greatest for stimuli that elicit weak bottom-up responses.

### Larger responses in IPS in high-demand tasks compared to low-demand tasks

Why does the brain disproportionately enhance responses to some stimuli and under some tasks compared to other experimental conditions? We suggest that this flexibility in attentional enhancement reflects the interaction between attention and the process of evidence accumulation to accomplish a perceptual decision. In this sense, attention is just one component of a perceptual task and we must consider other top-down processes, such as decision-making, when interpreting top-down modulation of neural responses. Stimuli with certain properties (e.g., low contrast, low phase-coherence) may yield weak or noisy sensory signals, and may therefore require extra decision time to complete the evidence accumulation process. We have identified IPS as a potential region that forms perceptual decisions, and this was evidenced by the fact that an evidence-accumulation model can link behavioral reaction times and IPS activity (Kay KN and JD Yeatman 2017).

Following this approach, we compared IPS responses across the various stimulus manipulations and tasks. IPS exhibited greater activity in the face task compared with the dot task in the position study (Figure 8A-B). This is in line with the more pronounced attentional effects observed in VTC for the face task. The result also mirrors the finding of greater IPS activity in the one-back task compared to the categorization task in the category study (Figure 8C-D). The flexible-attention framework also suggests that IPS activity may show systematic variation as a function of stimuli within a given task. Indeed, the correlations between IPS activity to the contrast and phase-coherence levels were established in (Kay KN and JD Yeatman 2017). However, in the position study, we did not find systematic correlation between IPS activity and attentional effects as function of eccentricity, possibly due to the limited slice coverage and suboptimal experimental design (see Discussion).

**Figure 8.**
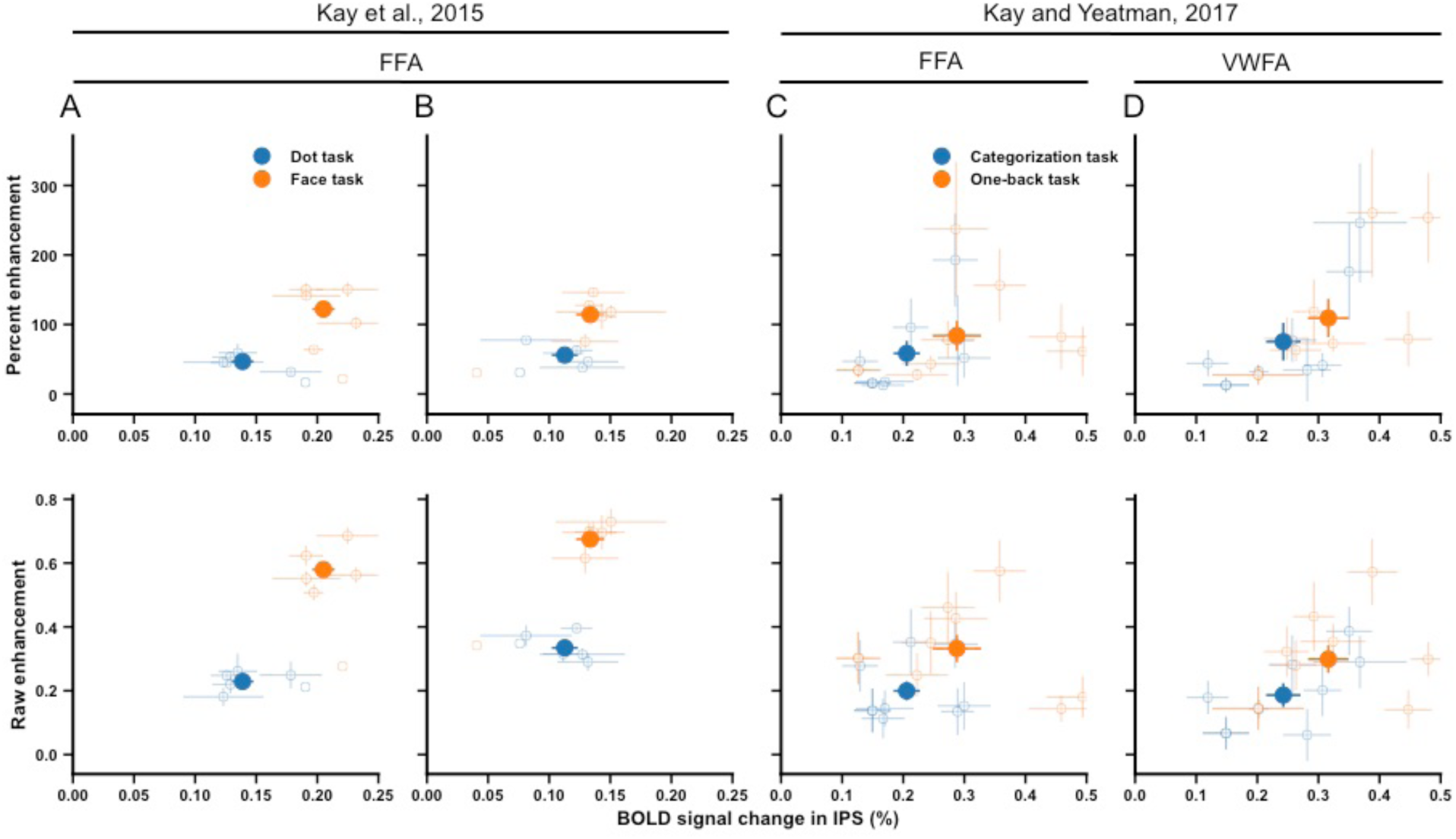
Percent enhancement and raw enhancement as a function of IPS activity. ***A-B***. Data from the position study (task experiment and interleaved-task experiment, respectively). The small open dots indicate different eccentricities, and the large solid dot indicates the mean across all locations. ***C-D***. Data from the category study. The small open dots indicate individual contrast and phase-coherence levels, and the large solid dot indicates the overall mean. The results show that IPS activity is larger for the face task compared to the dot task and for the one-back task compared to the categorization task, and this is accompanied by larger attentional enhancements in FFA and VWFA.

## DISCUSSION

In this article, we analyzed cortical responses in human VTC as a function of stimulus eccentricity and task. We found that the degree of attention-induced response enhancement increases from fovea to periphery and from a face-unrelated task to a face-related task. Moreover, analyses revealed consistent results in an independent experiment in the same study and another study involving additional stimulus manipulations and ROIs. Taken together, these results provide new evidence for constraining theoretical models of attention, and suggest that the effects of attention are dependent on stimuli and task in ways that are not captured by simple parametric models of attention that have been previously proposed. Understanding the mechanisms of attention might require further delineating the interaction between attention and other cognitive processes (e.g., decision-making).

### Previous models of attention do not account for the observed effects

Most prior research on the quantitative nature of attention has investigated the impact of attention on the shape of CRFs (Li X *et al.* 2008; Murray SO 2008; Boynton GM 2009). This approach has prompted several influential computational frameworks, such as the response-gain model (McAdams CJ and JH Maunsell 1999), the contrast-gain model (Reynolds JH *et al.* 2000; Martinez-Trujillo JC and S Treue 2004), and the additive-shift model (Buracas GT and GM Boynton 2007). One attraction of this fixed-parameter approach is that data from monkey electrophysiological, human fMRI, and psychophysical studies can be analyzed and compared within a common mathematical framework. Boynton GM (2009) used CRF modeling to summarize findings from seven different studies. Among three fMRI studies in his analysis, results in Buracas GT and GM Boynton (2007) and Murray SO (2008) are better explained by the additive-shift model, while results in Li X *et al.* (2008) are better explained by the contrast-gain model. These models, however, do not provide satisfactory explanations for our data (see Results).

With regard to the contrast-gain model, it is theoretically possible that attention shifts contrast response functions very far to the left so that only the upper asymptotic part of responses are observed, and this might be one way of attempting to reconcile the contrast-gain model with our measurements. However, notice that the contrast-gain model predicts that attention should produce no response difference at high contrast (i.e., 100%), but we can still see clear response differences at 100% contrast as well as 100% phase coherence and at the fovea (upper row in Figure 6).

Another limitation of the CRF modeling approach is that it is essentially a descriptive approach that merely summarizes the apparent structure of data into a function with a few parameters. The approach does not attempt to characterize the neural source of attentional modulations, such as where and how top-down influences are generated. In contrast, our efforts to characterize the IPS as the source of top-down modulations provides an opportunity to study more directly the causes that underlie modulations of sensory responses.

### The flexible-attention framework takes into account stimulus and task

Cognitive tasks are remarkably diverse, imposing different task demands on neural processing. For example, the categorization task in the category study requires attention to the stimuli and decisions made upon them; the one-back task in the category study requires both attention and temporal maintenance of information. We propose a flexible-attention framework that postulates that attention enhances responses in task-relevant regions in order to process specific stimuli and meet certain task demands. We emphasize that this is a *framework* that implies a change of conceptual stance, as opposed to a fully quantitative model of attention. In this framework, the observed top-down modulations in an experiment — which might be conventionally referred to as “attention” — depend on the details of the other cognitive processes used to fulfill the task (e.g., decision-making, memory). Conventional fixed-parameter modeling approaches do not take these complexities into account. For example, even though attention can be allocated to two different stimuli in seemingly the same way, the task difficulty might differ for these stimuli and lead to differing neural effects (Ress D et al. 2000; Kay KN and JD Yeatman 2017).

Our results have shown the significance of the flexible-attention framework. The inverse relationship that we have demonstrated between the strength of bottom-up responses and the magnitude of attentional enhancement has a clear interpretation in the context of evidence-accumulation models of perceptual decision-making. Most visual tasks require the brain to accumulate sensory evidence to make a decision, and in general we may suppose that weak neural responses constitute weak sensory evidence, therefore leading to longer evidence-accumulation.

Note that the flexible-attention framework does not imply that weak neural responses *always* receive disproportionately large top-down modulation. If a task involves no demand for processing weak stimuli, the attentional effect on weak stimuli might be small. For an illustration, consider the fact that attentional effects are relatively small for 0% phase-coherence stimuli (Figure 6D-F). It may be the case that the absence of any coherent form in these stimuli may render perceptual decisions (such as category judgment or one-back judgments) easier compared to the case of partially coherent stimuli. Accordingly, the evidence-accumulation process may be quite short. To more definitively resolve these unknowns, it is necessary to develop formal characterizations of the processes that underlie different tasks.

### IPS as a potential source of top-down attentional enhancement

One might wonder whether the flexible-attention framework can be translated into a quantitative model that characterizes neural responses. We proposed one such model, called IPS-scaling model, in the category study (Kay KN and JD Yeatman 2017). Researchers have long highlighted the crucial role of the parietal cortex in top-down attentional control; yet quantitative models have been rarely established. One important stride we made in the category study is to show that IPS activity predicts the amount of task-induced response scaling observed in FFA and VWFA.

We extend this analysis to the data from the position study. As shown in Figure 8A-B, IPS responses increase from the dot task to the face task, which mirrors the increase in top-down modulation in VTC from the dot task to the face task. However, we did not find systematic IPS-attention covariation across stimulus eccentricities within a task. This is possibly due to the specific experimental setting here. First, the position study did not set out to study interactions between IPS and VTC, and the scanning protocol provided only limited coverage of IPS (approximately up to IPS-0). This may have contributed to the noisy measurements of IPS responses (large horizontal error bars in Figure 8A-B). Second, the experimental design of the position study might not have been optimal for eliciting strong responses from the IPS. This is because the very quick presentation of stimuli (500ms/face) forces participants to quickly make decisions and this may preclude the complete unfolding of an evidence-accumulation process.

### Stronger attentional effect in high-level visual areas

In the present study, we primarily focused on high-level category-selective visual regions instead of low-level or middle-level visual regions, which are the focus of previous studies. One benefit of choosing FFA and VWFA is that we have relatively advanced understandings of their functional selectivities (Grill-Spector K et al. 2017). Moreover, these high-level visual areas are known for receiving greater attentional impacts compared to low-level visual areas (Kastner S and LG Ungerleider 2000). Indeed, we found much stronger attentional effects in high-level face-selective areas than low-level areas (Figures 3-4). This provides the advantage of a larger dynamic range of attentional enhancement, which helps to adjudicate different models of attention.

Another departure from previous studies is that we not only target high-level visual areas, but also measure their responses to a wide range of stimulus and task manipulations. For instance, previous studies using the CRF modeling approach typically manipulate only stimulus contrast. How attention influences visual coding on a broader range of feature dimensions (e.g., eccentricity, phase coherence) remains under-studied. In the position and category studies, we probed attentional effects as a function of three stimulus features (eccentricity, contrast, phase coherence), providing a more complete characterization of functional properties of the visual system. One recent study found that attentional effects were larger in fovea than in periphery (Bressler DW et al. 2013). That study, however, used checkerboard stimuli that elicited only strong responses in low-level visual areas. One important direction of future work might be to explain such differential attentional effect between low-level and high-level cortices.

### Region-level characterization of attentional effects

The original analyses performed in the position study (Kay KN *et al.* 2015) examined attentional effects on spatial representation in human VTC at the level of single voxels. Through population receptive field (pRF) modeling, it was shown that task-specific attention alters the center, size and amplitude of pRFs of voxels in VTC. Analyses in the current paper pursue a fundamentally different approach and provide different insight into the nature of attentional modulation. Rather than characterizing the spatial tuning profile of individual voxels, we calculated region-level responses and investigated how and why the strength of attentional modulations varies for different stimuli and tasks. Though the motivations are different, the two approaches have revealed some conceptually consistent results. For instance, both analyses demonstrate greater attentional effects in the face task compared to the dot task, and attentional effects are found to progressively develop along the visual hierarchy. We believe that exploring different analyses and interpretations of neural measurements is critical for achieving a better understanding of attentional effects.

## AUTHOR CONTRIBUTIONS

R.Z. analyzed the data. R.Z. and K.K. wrote the paper.

## ACKNOWLEDGEMENTS

We thank S. Engel for comments on the manuscript. This work was supported by NIH Grants P41 EB015894 and P30 NS076408.

